## Supplementary Material for "Neutral vs. non-neutral genetic footprints of *Plasmodium falciparum* multiclonal infections"

### Figures

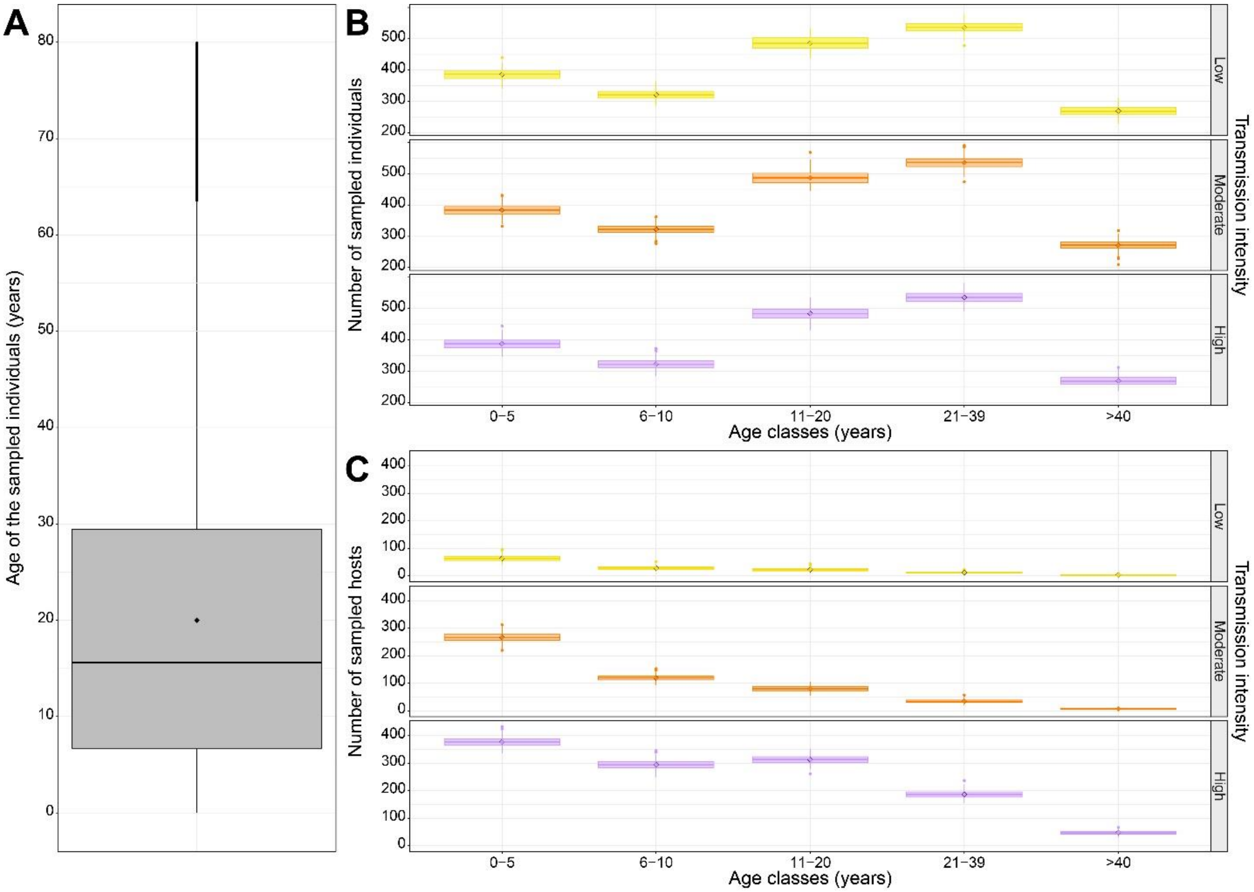


**S1 Fig. Host age distribution, and number of sampled individuals and hosts per age class.** For each category, the horizontal central solid line represents the median, the diamond represents the mean, the box represents the interquartile range (IQR) from the 25th to 75th centiles, the whiskers indicate the most extreme data point which is no more than 1.5 times the interquartile range from the box, and the dots show the outliers, i.e. the points beyond the whiskers. **A)** Age distribution of the sampled individuals. **B)** Number of sampled individuals per age class. **C)** Number of sampled hosts per age class. Upper (yellow), middle (orange), and lower (purple) panels correspond to simulations under low-, moderate-, and high-transmission settings, respectively (S1 and S2 Tables). Values were split into five age classes, i.e. 0-5, 6-10, 11-20, 21-39, and ≥ 40 years.

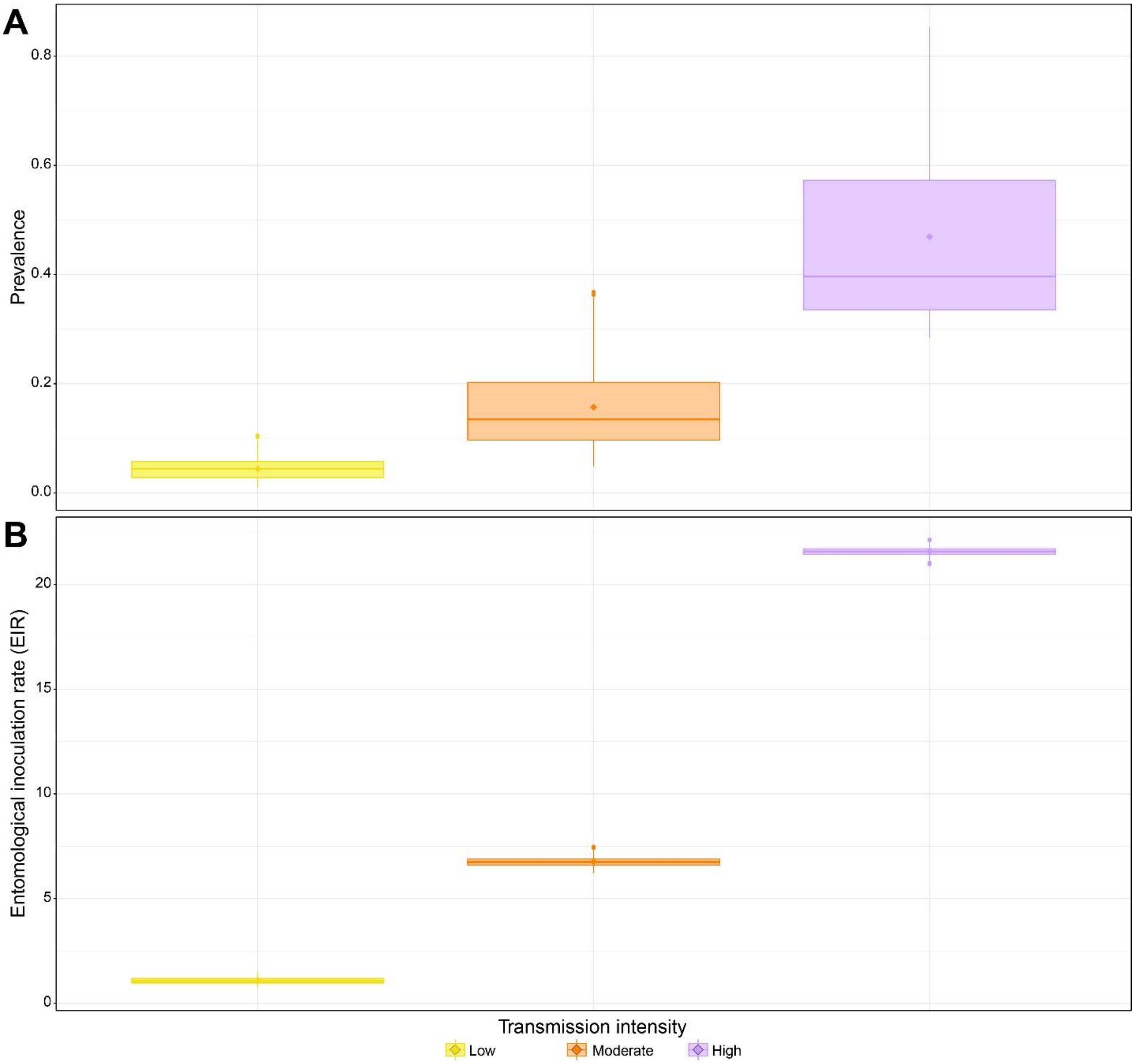


**S2 Fig. Prevalence and entomological inoculation rate (EIR) per transmission intensity.** For each category, the horizontal central solid line represents the median, the diamond represents the mean, the box represents the interquartile range (IQR) from the 25th to 75th centiles, the whiskers indicate the most extreme data point which is no more than 1.5 times the interquartile range from the box, and the dots show the outliers, i.e. the points beyond the whiskers. **A)** Prevalence; **B)** EIR. Statistics calculated for simulations under low-, moderate-, and high-transmission settings are indicated in yellow, orange, and purple, respectively (S1 and S2 Tables).

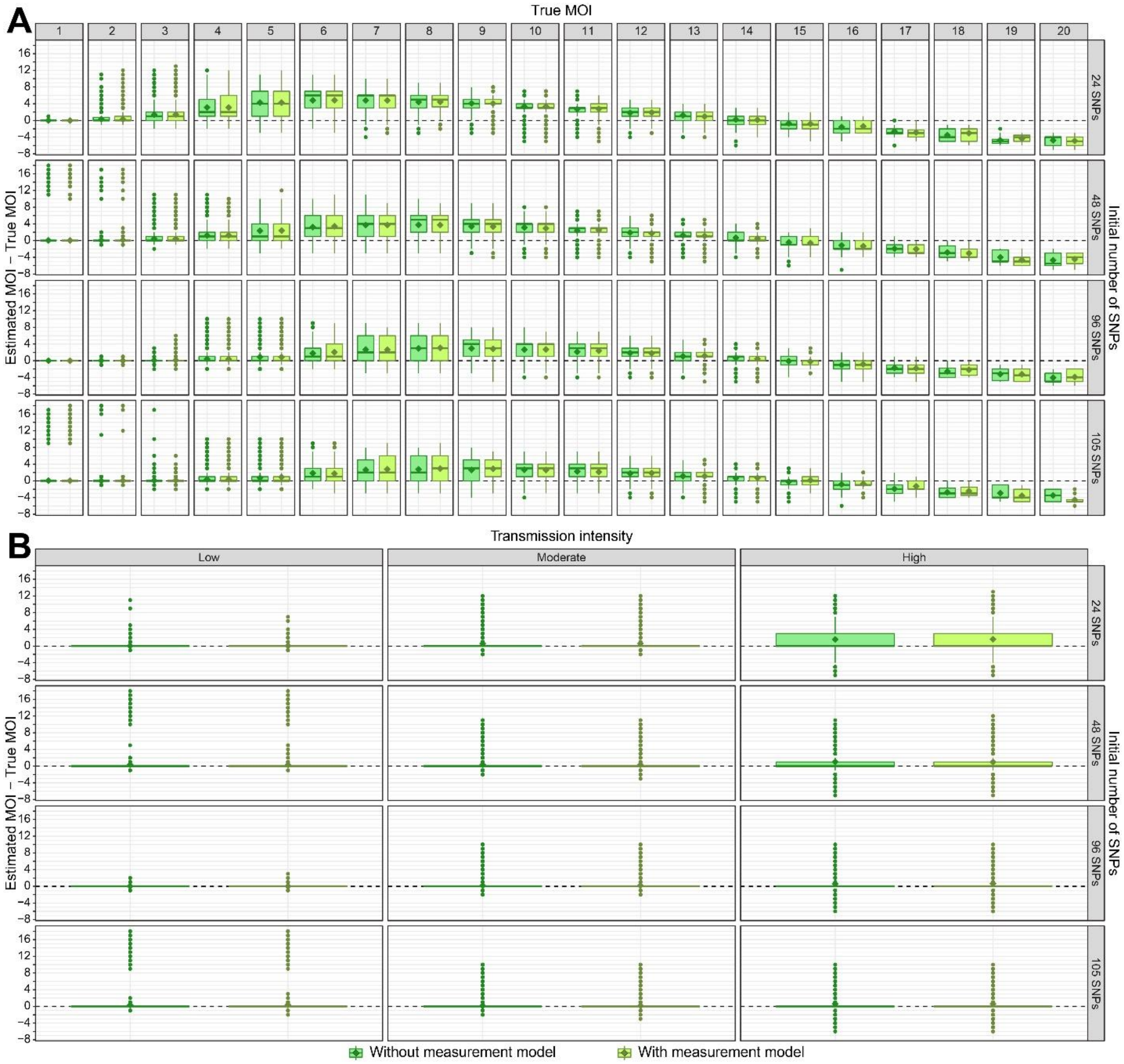


**S3 Fig. Initial number of SNPs and accuracy of the multiplicity of infection (MOI) estimates determined with THE REAL McCOIL approach.** The accuracy of MOI estimates is defined as the differences between estimated and true MOI per host. While null values highlight accurate MOI estimates (indicated by a dashed black horizontal line), the positive and negative values highlight over- and under-estimation, respectively. The dark and light green colors indicate respectively MOI estimations made without and with a measurement model (Fig 2). For each category, the horizontal central solid line represents the median, the diamond represents the mean, the box represents the interquartile range (IQR) from the 25th to 75th centiles, the whiskers indicate the most extreme data point which is no more than 1.5 times the interquartile range from the box, and the dots show the outliers, i.e. the points beyond the whiskers. **A)** Accuracy of MOI estimates per true MOI. **B)** Accuracy of MOI estimates per transmission intensity (S1 and S2 Tables).

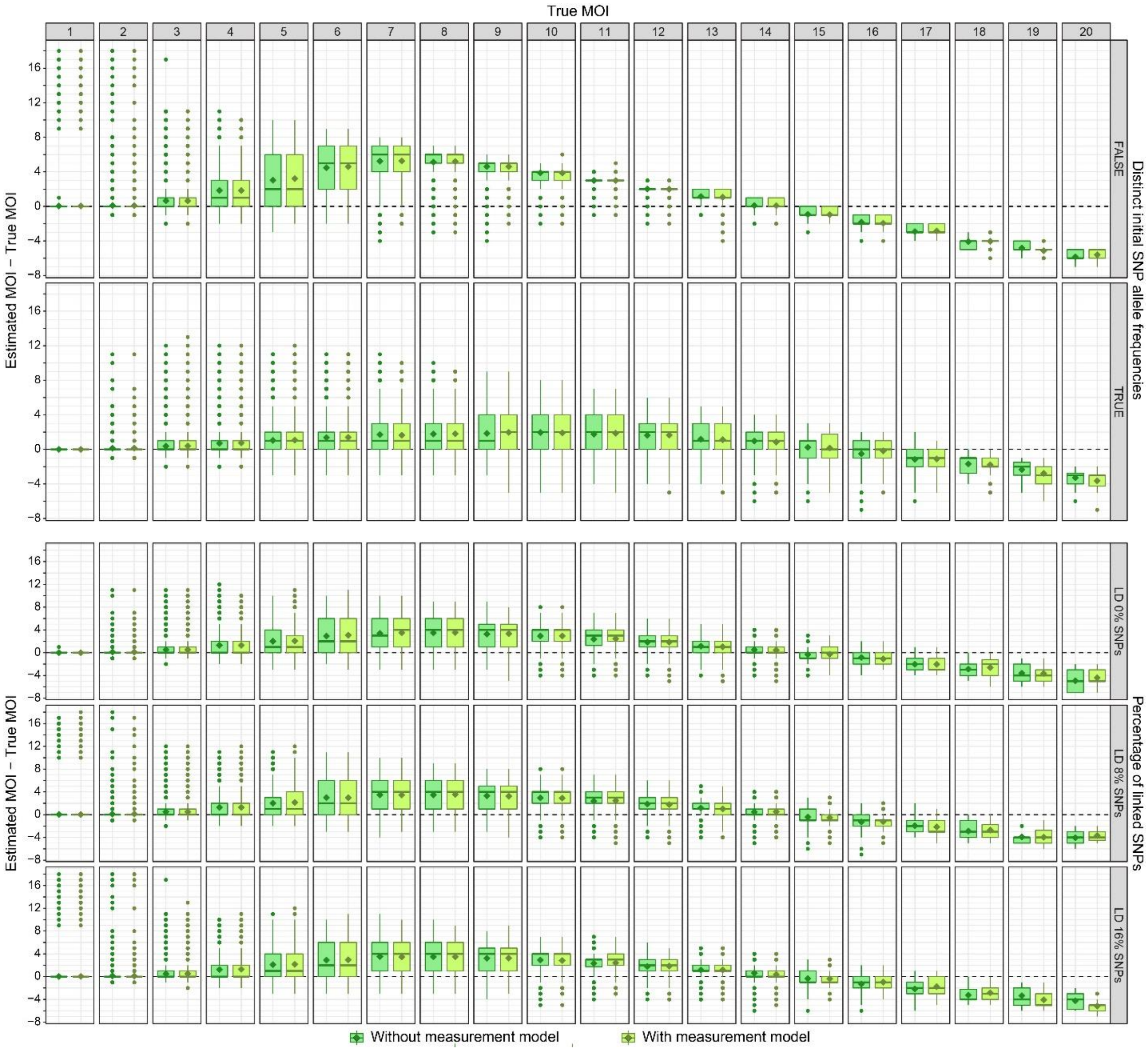


**S4 Fig. SNP properties and accuracy of the multiplicity of infection (MOI) estimates determined with THE REAL McCOIL approach.** The accuracy of MOI estimates is defined as the differences between estimated and true MOI per host. While null values highlight accurate MOI estimates (indicated by a dashed black horizontal line), the positive and negative values highlight over- and under-estimation, respectively. For each category, the horizontal central solid line represents the median, the diamond represents the mean, the box represents the interquartile range (IQR) from the 25th to 75th centiles, the whiskers indicate the most extreme data point which is no more than 1.5 times the interquartile range from the box, and the dots show the outliers, i.e. the points beyond the whiskers. The dark and light green colors indicate respectively MOI estimations made without and with a measurement model (Fig 2).

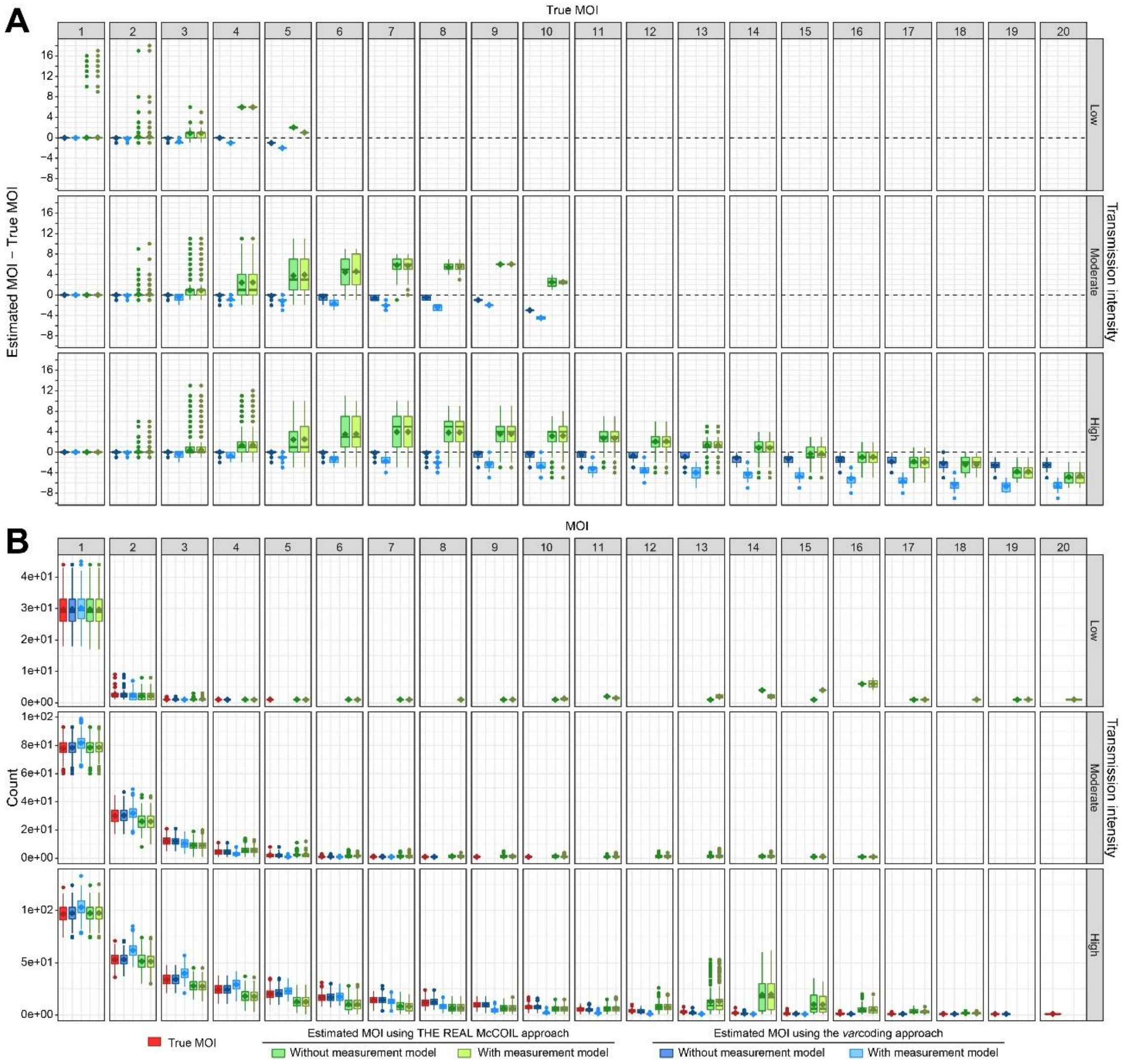


**S5 Fig: Reliability of the multiplicity of infection (MOI) estimations when subsampling 25% of the sampled individuals (i.e. 500 individuals).** For each category, the horizontal central solid line represents the median, the diamond represents the mean, the box represents the interquartile range (IQR) from the 25th to 75th centiles, the whiskers indicate the most extreme data point which is no more than 1.5 times the interquartile range from the box, and the dots show the outliers, i.e. the points beyond the whiskers. The upper, middle, and lower row panels correspond to simulations under low-, moderate-, and high-transmission settings, respectively (S1 and S2 Tables). **A)** Accuracy of MOI estimates, defined as the difference between estimated and true MOI per host. While null values highlight accurate MOI estimates (indicated by a dashed black horizontal line), the positive and negative values highlight over- and under-estimation, respectively. Estimates with the neutral SNP-based approach (THE REAL McCOIL) are indicated in green, and those with the *var* gene-based approach (*var*coding) are indicated in blue. The dark and light green or blue colors indicate respectively MOI estimations made without and with a measurement model (Fig 2). The column panels show differences for specific true MOI values. **B)** Population distribution of the estimated and true MOI per host from the simulated "true" values and those estimated with the methods indicated by the colors similar to panel A.

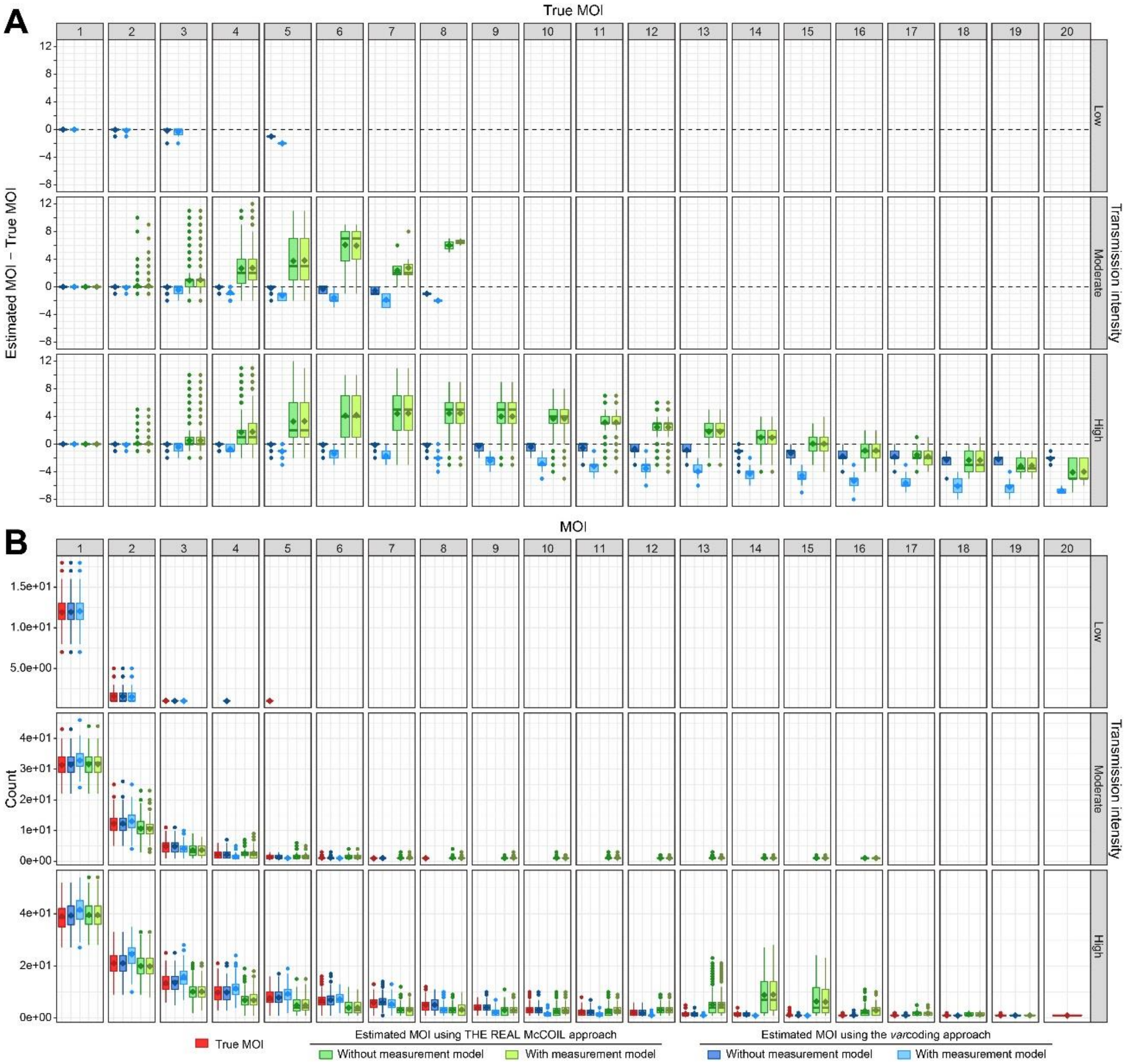


**S6 Fig: Reliability of the multiplicity of infection (MOI) estimations when subsampling 10% of the sampled individuals (i.e. 200 individuals).** For each category, the horizontal central solid line represents the median, the diamond represents the mean, the box represents the interquartile range (IQR) from the 25th to 75th centiles, the whiskers indicate the most extreme data point which is no more than 1.5 times the interquartile range from the box, and the dots show the outliers, i.e. the points beyond the whiskers. The upper, middle, and lower row panels correspond to simulations under low-, moderate-, and high-transmission settings, respectively (S1 and S2 Tables). **A)** Accuracy of MOI estimates, defined as the difference between estimated and true MOI per host. While null values highlight accurate MOI estimates (indicated by a dashed black horizontal line), the positive and negative values highlight over- and under-estimation, respectively. Estimates with the neutral SNP-based approach (THE REAL McCOIL) are indicated in green, and those with the *var* gene-based approach (*var*coding) are indicated in blue. The dark and light green or blue colors indicate respectively MOI estimations made without and with a measurement model (Fig 2). The column panels show differences for specific true MOI values. **B)** Population distribution of the estimated and true MOI per host from the simulated "true" values and those estimated with the methods indicated by the colors similar to panel A.

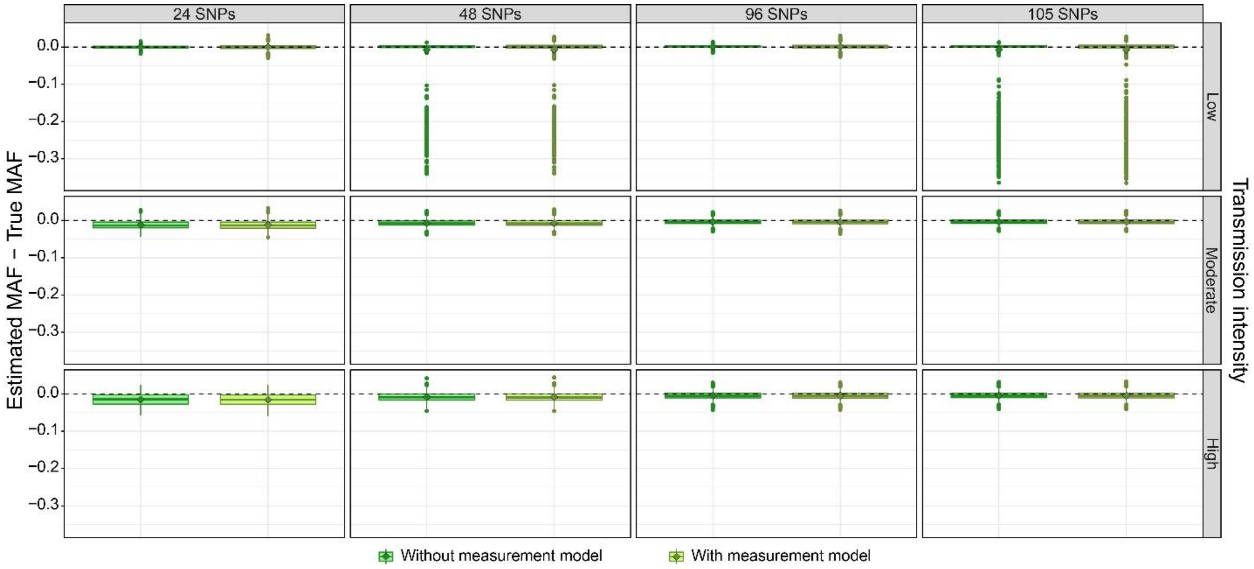


**S7 Fig. Accuracy of the minor allele frequency (MAF) estimates per locus determined with THE REAL McCOIL approach.** The accuracy of MAF estimates per locus is defined as the differences between estimated and true MAF per locus. While null values highlight accurate MAF estimates per locus (indicated by a dashed black horizontal line), the positive and negative values highlight over- and under-estimation, respectively. For each category, the horizontal central solid line represents the median, the diamond represents the mean, the box represents the interquartile range (IQR) from the 25th to 75th centiles, the whiskers indicate the most extreme data point which is no more than 1.5 times the interquartile range from the box, and the dots show the outliers, i.e. the points beyond the whiskers. The dark and light green colors indicate respectively MAF estimations made without and with a measurement model (Fig 2). Upper, middle, and lower panels correspond to simulations under low-, moderate-, and high-transmission settings, respectively (S1 and S2 Tables).

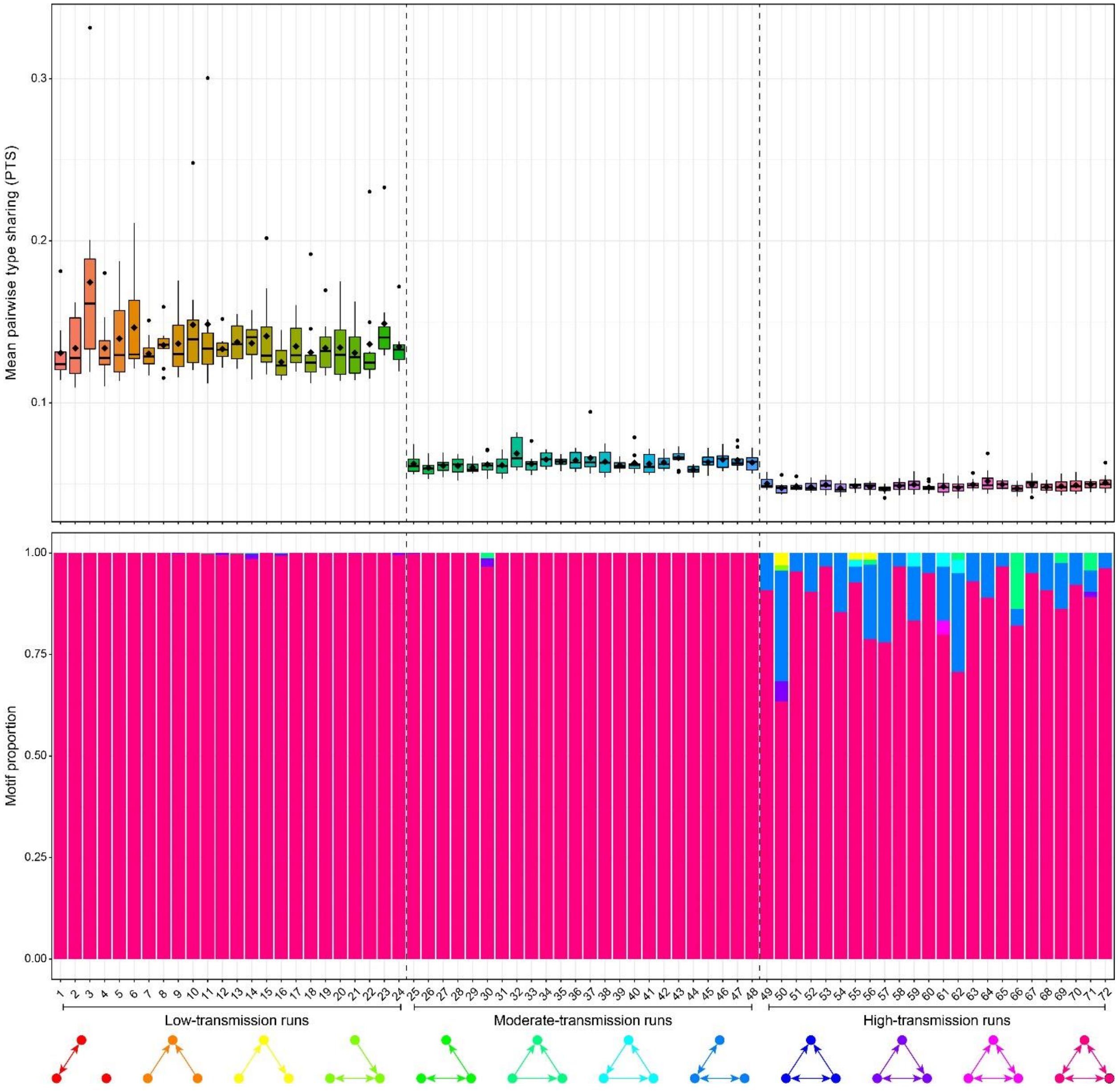


**S8 Fig. Population structure using repertoire similarity network properties.** Comparisons of repertoire similarity networks of 150 randomly sampled parasite *var* repertoires generated from a one-time point under low, moderate, and high-transmission settings (S1 and S2 Tables). Only the top 1% of edges are drawn and used in the analysis. The upper panel shows the distribution of the mean pairwise type sharing (PTS) per run. For each category, the horizontal central solid line represents the median, the diamond represents the mean, the box represents the interquartile range (IQR) from the 25th to 75th centiles, the whiskers indicate the most extreme data point which is no more than 1.5 times the interquartile range from the box, and the dots show the outliers, i.e. the points beyond the whiskers. The lower panel shows the distributions of the proportion of occurrences of three-node graph motifs across the repertoire similarity networks.

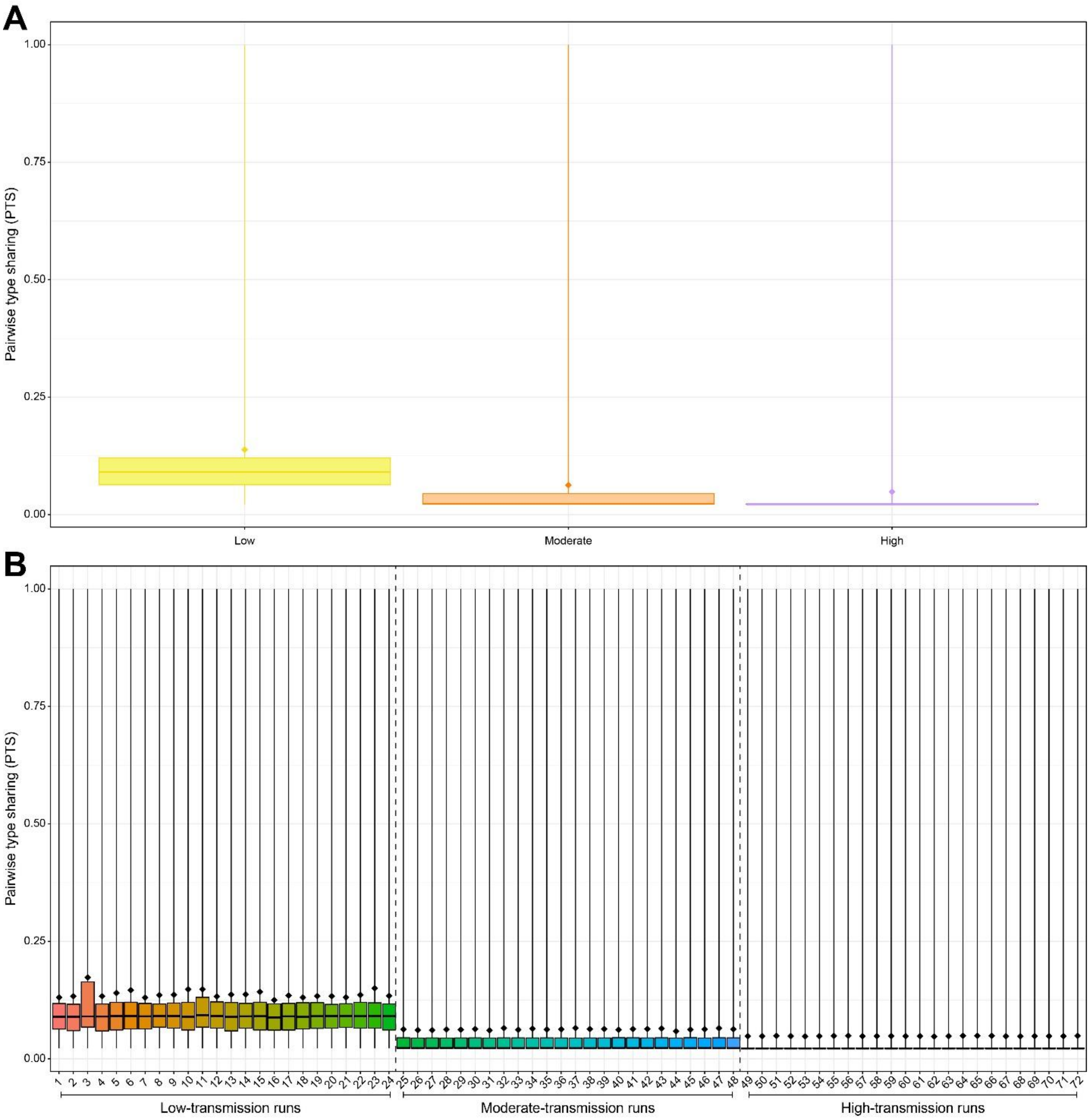


**S9 Fig. Pairwise type sharing (PTS).** For each category, the horizontal central solid line represents the median, the diamond represents the mean, the box represents the interquartile range (IQR) from the 25th to 75th centiles, and the whiskers indicate the most extreme data point. **A)** Distribution of the PTS per transmission intensity. **B)** Distribution of the PTS per run.

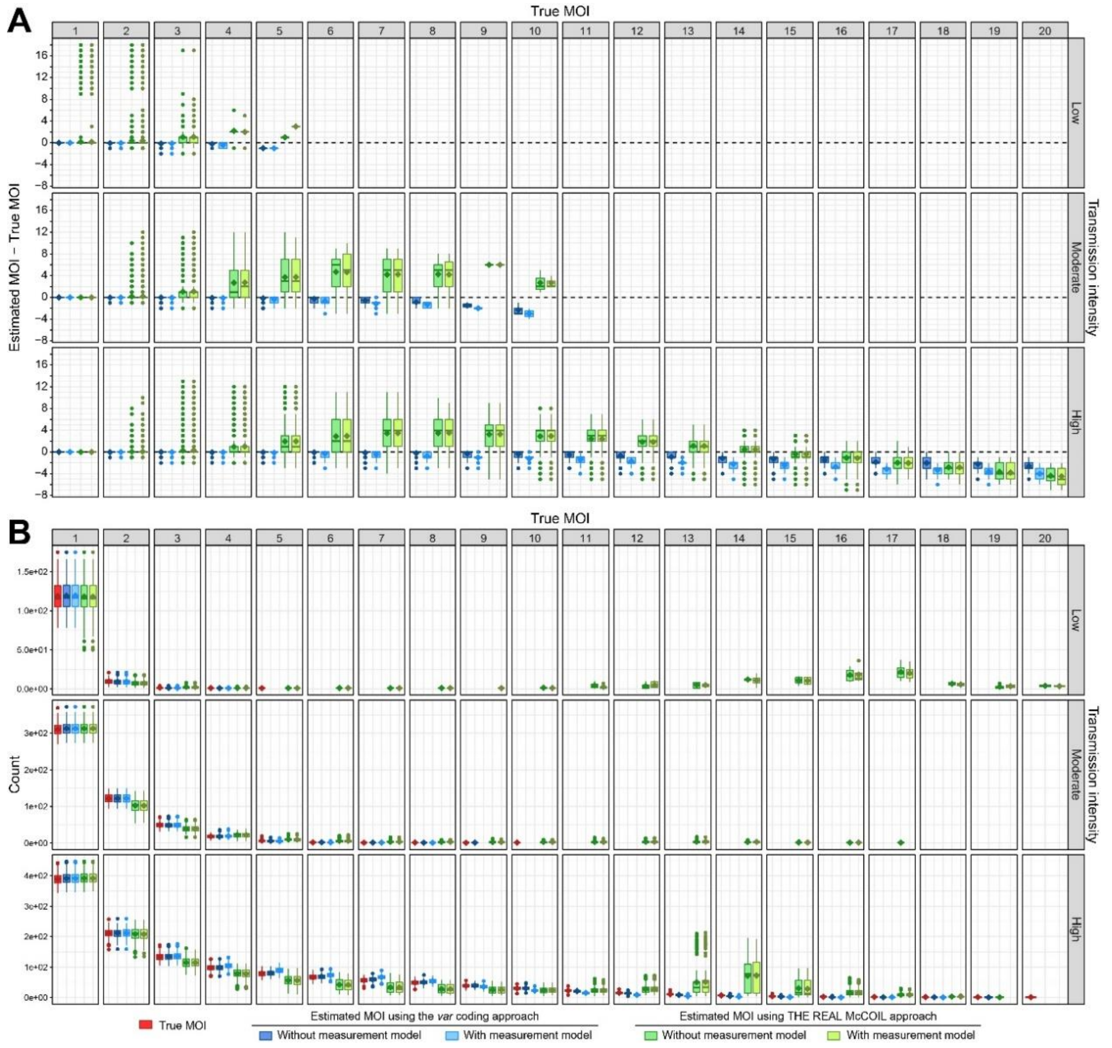


**S10 Fig. Reliability of the multiplicity of infection (MOI) estimations when simulations include a measurement error based on the distribution of the number of non-upsA DBLα *var* gene types per 3D7 laboratory isolates for the *var* coding approach.** For each category, the horizontal central solid line represents the median, the diamond represents the mean, the box represents the interquartile range (IQR) from the 25th to 75th centiles, the whiskers indicate the most extreme data point which is no more than 1.5 times the interquartile range from the box, and the dots show the outliers, i.e. the points beyond the whiskers. The upper, middle, and lower row panels correspond to simulations under low-, moderate-, and high-transmission settings, respectively (S1 and S2 Tables). **A)** Accuracy of MOI estimates, defined as the differences between estimated and true MOI per host. While null values highlight accurate MOI estimates (indicated by a dashed black horizontal line), the positive and negative values highlight over- and under-estimation, respectively. Estimates with the neutral SNP-based approach (THE REAL McCOIL) are indicated in green, and those with the *var* gene-based approach (*var*coding) are indicated in blue. The dark and light blue or green colors indicate respectively MOI estimates made without and with a measurement model (Fig 2). The column panels show differences for specific true MOI values. **B)** Population distribution of the estimated and true MOI per host from the simulated “true” values and those estimated with the methods indicated by the colors similar to panel A.

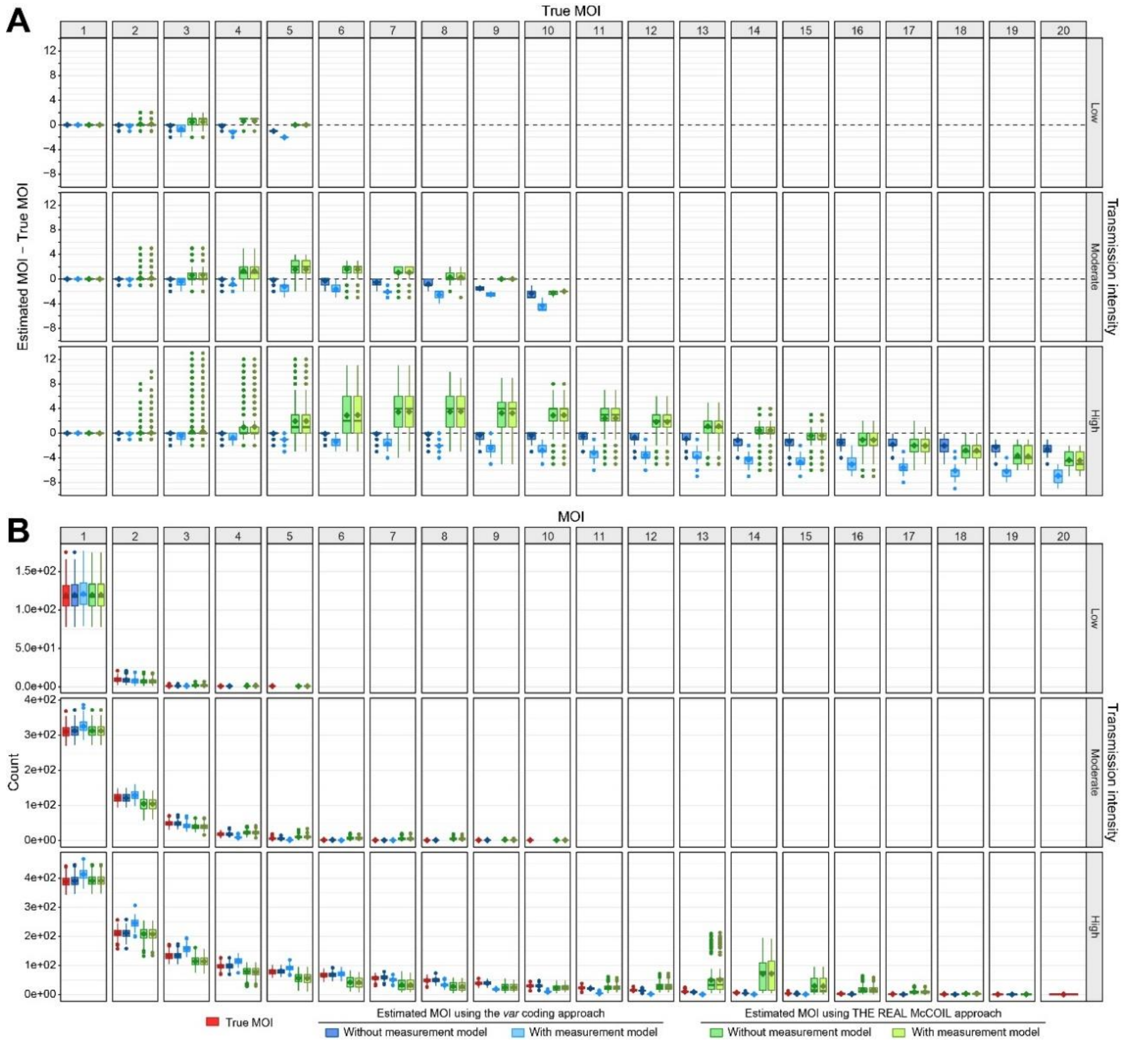


**S11 Fig. Reliability of the multiplicity of infection (MOI) estimations when THE REAL McCOIL approach using an upper bound for MOI of 5, 10, and 20 for the low-, moderate-, and high-transmission simulations, respectively.** For each category, the horizontal central solid line represents the median, the diamond represents the mean, the box represents the interquartile range (IQR) from the 25th to 75th centiles, the whiskers indicate the most extreme data point which is no more than 1.5 times the interquartile range from the box, and the dots show the outliers, i.e. the points beyond the whiskers. The upper, middle, and lower row panels correspond to simulations under low-, moderate-, and high-transmission settings, respectively (S1 and S2 Tables). **A)** Accuracy of MOI estimates, defined as the differences between estimated and true MOI per host. While null values highlight accurate MOI estimates (indicated by a dashed black horizontal line), the positive and negative values highlight over- and under-estimation, respectively. The estimated MOI using the *var* genes based approach (i.e. *var* coding) are indicated in blue, and the estimated MOI using the neutral SNPs based approach (i.e. THE REAL McCOIL) are indicated in green. The dark and light blue or green colors indicate respectively MOI estimates made without and with a measurement model (Fig 2). The column panels show differences for specific true MOI values. **B)** Population distribution of the estimated and true MOI per host from the simulated "true" values and those estimated with the methods indicated by the colors similar to panel A.

### Tables

**S1 Table.** Epidemiological and genetic parameters used in the stochastic simulations.

| Name | Values | Description |
|---|---|---|
| biting_rate | [5.0E-05, 7.5E-05, 1.0E-04] * daily_biting_rate_multiplier | Transmission rate for each day of the year. |
| coinfection_reduces_transmission | true | Whether or not transmissibility is reduced with coinfection. |
| distinct_initial_snp_allele_frequencies | [true, false] | Whether the initial allele frequencies of the SNPs are distinct. |
| ectopic_recombination_generates_new_alleles | false | Whether or not ectopic recombination generates new alleles. |
| ectopic_recombination_rate | 1.80E-07 | Ectopic recombination rate parameter. |
| gene_strain_count_period | 30 | How often to output the number of circulating genes and strains. |
| host_sample_size | 2000 | Number of hosts to sample at each sampling period. |
| host_sampling_period | 30 | How often to sample host output. |
| immigration_rate_fraction | 0.0026 | Immigration rate, as a fraction of the non-immigration biting rate. |
| immunity_level_max | 100 | Maximum immunity level. |
| immunity_loss_rate | 0.001 | Rate at which immunity is lost, per host, per gene. |
| initial_snp_allele_frequency | [0.1 0.9] | Range of the possible initial frequencies for one of the two SNP alleles. |
| max_host_lifetime | 80 * t_year | Maximum host lifetime. |
| mean_host_lifetime | 30 * t_year | Mean of exponential distribution used to draw host lifetime. |
| mean_n_mutations_per_epitope | 5 | Mean number of mutations per epitope for similarity calculation. |
| migrants_match_local_prevalence | true | Whether the immigration rate needs to time the local infection rate. |
| mutation_rate | 1.42E-08 | Rate of mutation, per active infection. |
| n_alleles_per_locus_initial | n_genes_initial / 10 | Initial number of alleles for each epitope locus. |
| n_genes_initial | [500, 2000, 10000] | Number of genes in the initial gene pool. |
| n_genes_per_strain | 45 | Number of genes in strain. |
| n_hosts | 10000 | Number of hosts. |
| n_infections_active_max | 20 | Maximum number of simultaneous active infections. |
| n_infections_liver_max | 20 | Maximum number of simultaneous infections in the liver stage. |
| n_initial_infections | 20 | Number of initial infections. |
| n_loci | 2 | Number of epitope loci in each gene. |
| n_snps_per_strain | [24, 48, 96, and 105] | Number of biallelic neutral single nucleotide polymorphims (SNPs) in strain. |
| p_ectopic_recombination_is_conversion | 0 | Probability that an ectopic recombination is a conversion. |
| rho_recombination_tolerance | 0.8 | Recombination tolerance, rho, Drummond et al. |
| rng_seed | nothing | Seed for random number generator. |
| sample_duration | 1000 | Sample an infection duration every `sample_duration` infection(s). |
| snp_linkage_disequilibrium | [true, false] | Whether the SNPs (or some SNPs) are in linkage disequilibrium (LD). |
| snp_pairwise_ld | snp_ld_matrix | Pairwise linkage disequilibrium (LD) matrix. |
| summary_period | 30 | How often to write summary output. |
| switching_rate | 1.0/6.0 | Switching rate for genes the host is not immune to. |
| transmissibility | 0.5 | Baseline transmissibility of infections. |
| t_liver_stage | 14 | Duration of the liver stage. |
| t_burnin | 30240 | Burn-in time. |
| t_end | 30600 | Simulation end time. |
| t_year | 360 | Number of time units in a year. |
| upper_bound_recomputation_period | 30 | How often to recompute upper bounds for rejection sampling. |
| use_immunity_by_allele | true | Immunity model. |
| verification_period | 30 | How often to verify consistency of simulation state. |
| whole_gene_immune | false | Whether a host gains immunity towards a gene if the host has seen all the alleles. |

**S2 Table.** Epidemiological and genetic distinct parameters per run.

| Run | n_genes_ initial | n_snps_per strain | biting_rate_mean | distinct_initial_snp _allele_frequencies | snp_linkage_ disequilibrium | snp_pairwise_ld |
|---|---|---|---|---|---|---|
| 1 | 500 | 24 | 5.00E-05 | true | true | 8% linked SNPs |
| 2 | 500 | 24 | 5.00E-05 | true | true | 16% linked SNPs |
| 3 | 500 | 24 | 5.00E-05 | true | false | 0% linked SNPs |
| 4 | 500 | 24 | 5.00E-05 | false | true | 8% linked SNPs |
| 5 | 500 | 24 | 5.00E-05 | false | true | 16% linked SNPs |
| 6 | 500 | 24 | 5.00E-05 | false | false | 0% linked SNPs |
| 7 | 500 | 48 | 5.00E-05 | true | true | 8% linked SNPs |
| 8 | 500 | 48 | 5.00E-05 | true | true | 16% linked SNPs |
| 9 | 500 | 48 | 5.00E-05 | true | false | 0% linked SNPs |
| 10 | 500 | 48 | 5.00E-05 | false | true | 8% linked SNPs |
| 11 | 500 | 48 | 5.00E-05 | false | true | 16% linked SNPs |
| 12 | 500 | 48 | 5.00E-05 | false | false | 0% linked SNPs |
| 13 | 500 | 96 | 5.00E-05 | true | true | 8% linked SNPs |
| 14 | 500 | 96 | 5.00E-05 | true | true | 16% linked SNPs |
| 15 | 500 | 96 | 5.00E-05 | true | false | 0% linked SNPs |
| 16 | 500 | 96 | 5.00E-05 | false | true | 8% linked SNPs |
| 17 | 500 | 96 | 5.00E-05 | false | true | 16% linked SNPs |
| 18 | 500 | 96 | 5.00E-05 | false | false | 0% linked SNPs |
| 19 | 500 | 105 | 5.00E-05 | true | true | 8% linked SNPs |
| 20 | 500 | 105 | 5.00E-05 | true | true | 16% linked SNPs |
| 21 | 500 | 105 | 5.00E-05 | true | false | 0% linked SNPs |
| 22 | 500 | 105 | 5.00E-05 | false | true | 8% linked SNPs |
| 23 | 500 | 105 | 5.00E-05 | false | true | 16% linked SNPs |
| 24 | 500 | 105 | 5.00E-05 | false | false | 0% linked SNPs |
| 25 | 2,000 | 24 | 7.50E-05 | true | true | 8% linked SNPs |
| 26 | 2,000 | 24 | 7.50E-05 | true | true | 16% linked SNPs |
| 27 | 2,000 | 24 | 7.50E-05 | true | false | 0% linked SNPs |
| 28 | 2,000 | 24 | 7.50E-05 | false | true | 8% linked SNPs |
| 29 | 2,000 | 24 | 7.50E-05 | false | true | 16% linked SNPs |
| 30 | 2,000 | 24 | 7.50E-05 | false | false | 0% linked SNPs |
| 31 | 2,000 | 48 | 7.50E-05 | true | true | 8% linked SNPs |
| 32 | 2,000 | 48 | 7.50E-05 | true | true | 16% linked SNPs |
| 33 | 2,000 | 48 | 7.50E-05 | true | false | 0% linked SNPs |
| 34 | 2,000 | 48 | 7.50E-05 | false | true | 8% linked SNPs |
| 35 | 2,000 | 48 | 7.50E-05 | false | true | 16% linked SNPs |
| 36 | 2,000 | 48 | 7.50E-05 | false | false | 0% linked SNPs |
| 37 | 2,000 | 96 | 7.50E-05 | true | true | 8% linked SNPs |
| 38 | 2,000 | 96 | 7.50E-05 | true | true | 16% linked SNPs |
| 39 | 2,000 | 96 | 7.50E-05 | true | false | 0% linked SNPs |
| 40 | 2,000 | 96 | 7.50E-05 | false | true | 8% linked SNPs |
| 41 | 2,000 | 96 | 7.50E-05 | false | true | 16% linked SNPs |
| 42 | 2,000 | 96 | 7.50E-05 | false | false | 0% linked SNPs |
| 43 | 2,000 | 105 | 7.50E-05 | true | true | 8% linked SNPs |
| 44 | 2,000 | 105 | 7.50E-05 | true | true | 16% linked SNPs |
| 45 | 2,000 | 105 | 7.50E-05 | true | false | 0% linked SNPs |
| 46 | 2,000 | 105 | 7.50E-05 | false | true | 8% linked SNPs |
| 47 | 2,000 | 105 | 7.50E-05 | false | true | 16% linked SNPs |
| 48 | 2,000 | 105 | 7.50E-05 | false | false | 0% linked SNPs |
| 49 | 10,000 | 24 | 1.00E-04 | true | true | 8% linked SNPs |
| 50 | 10,000 | 24 | 1.00E-04 | true | true | 16% linked SNPs |
| 51 | 10,000 | 24 | 1.00E-04 | true | false | 0% linked SNPs |
| 52 | 10,000 | 24 | 1.00E-04 | false | true | 8% linked SNPs |
| 53 | 10,000 | 24 | 1.00E-04 | false | true | 16% linked SNPs |
| 54 | 10,000 | 24 | 1.00E-04 | false | false | 0% linked SNPs |
| 55 | 10,000 | 48 | 1.00E-04 | true | true | 8% linked SNPs |
| 56 | 10,000 | 48 | 1.00E-04 | true | true | 16% linked SNPs |
| 57 | 10,000 | 48 | 1.00E-04 | true | false | 0% linked SNPs |
| 58 | 10,000 | 48 | 1.00E-04 | false | true | 8% linked SNPs |
| 59 | 10,000 | 48 | 1.00E-04 | false | true | 16% linked SNPs |
| 60 | 10,000 | 48 | 1.00E-04 | false | false | 0% linked SNPs |
| 61 | 10,000 | 96 | 1.00E-04 | true | true | 8% linked SNPs |
| 62 | 10,000 | 96 | 1.00E-04 | true | true | 16% linked SNPs |
| 63 | 10,000 | 96 | 1.00E-04 | true | false | 0% linked SNPs |
| 64 | 10,000 | 96 | 1.00E-04 | false | true | 8% linked SNPs |
| 65 | 10,000 | 96 | 1.00E-04 | false | true | 16% linked SNPs |
| 66 | 10,000 | 96 | 1.00E-04 | false | false | 0% linked SNPs |
| 67 | 10,000 | 105 | 1.00E-04 | true | true | 8% linked SNPs |
| 68 | 10,000 | 105 | 1.00E-04 | true | true | 16% linked SNPs |
| 69 | 10,000 | 105 | 1.00E-04 | true | false | 0% linked SNPs |
| 70 | 10,000 | 105 | 1.00E-04 | false | true | 8% linked SNPs |
| 71 | 10,000 | 105 | 1.00E-04 | false | true | 16% linked SNPs |
| 72 | 10,000 | 105 | 1.00E-04 | false | false | 0% linked SNPs |

**S3 Table.** Pearson correlation coefficients between the inaccuracy of the minor allele frequency (MAF) per locus estimated with THE REAL McCOIL approach (defined as the absolute differences between estimated and true MAF per locus), and the locus properties.

| **Locus properties** | | **Inaccuracy of the MAF estimates** | |
|---|---|---|---|
| | | **With measurement model** | **Without measurement model** |
| Proportion of double allele calls (DACs) | | -0.08 *** | -0.09 *** |
| Proportion of single allele calls | Major allele | 0.02 *** | 0.02 *** |
| | Minor allele | -0.05 *** | -0.08 *** |
| Proportion of missing allele calls | | 0.11 *** | 0.11 *** |
| True MAF | | -0.05 *** | -0.06 *** |

*** *P*-value < 0.001
